# Reinvestigating the role of ERK5: kinase activity is not required for cellular immune response or proliferation

**DOI:** 10.1101/038513

**Authors:** Emme C.K. Lin, Christopher M. Amantea, Tyzoon K. Nomanbhoy, Helge Weissig, Junichi Ishiyama, Yi Hu, Shyama Sidique, Bei Li, John W. Kozarich, Jonathan S. Rosenblum

**Affiliations:** ActivX Biosciences, Inc., 11025 N. Torrey Pines Rd, La Jolla, CA, 92037 United States of America; Watarase Research Center, Kyorin Pharmaceutical Co., Ltd., 1848 Nogi, Nogi-machi, Shimotsuga-gun, Tochigi 329-0114, Japan

**Keywords:** ERK5, bromodomain, inflammation, proliferation, kinase

## Abstract

Unlike other members of the MAPK family, ERK5 contains a large C-terminal domain with transcriptional activation capability in addition to an N-terminal canonical kinase domain. Genetic deletion of ERK5 is embryonic lethal and tissue-restricted deletions have profound effects on erythroid development, cardiac function and neurogenesis. In addition, depletion of ERK5 is anti-inflammatory and anti-tumorigenic. Small molecule inhibition of ERK5 has been shown to have promising activity in cell and animal models of inflammation and oncology. Here we report the synthesis and biological characterization of potent, selective ERK5 inhibitors. In contrast to both genetic depletion/deletion of ERK5 and inhibition with previously reported compounds, inhibition of the kinase with the most selective of the new inhibitors had no anti-inflammatory or anti-proliferative activity. The source of efficacy in previously reported ERK5 inhibitors is shown to be off-target activity on bromodomains (BRDs), conserved protein modules involved in recognition of acetyl-lysine residues during transcriptional processes. It is likely that phenotypes reported from genetic deletion or depletion of ERK5 arise from removal of a non-catalytic function of ERK5. The newly reported inhibitors should be useful in determining which of the many reported phenotypes are due to kinase activity, and delineate which can be pharmacologically targeted.

## Significance

Whole protein deletion and pharmacological inhibition are frequently used to functionally annotate enzymes. Each has limitations: whole protein deletion removes both enzymatic and non-enzymatic functions; small molecule inhibitors can have unrecognized off-target activities. When both approaches agree, it’s nearly incontrovertible support for protein function. Here we describe a counterexample. ERK5 knockdown and inhibition have supported a role for this kinase in a number of biological processes. We show that previously reported ERK5 compounds inhibit bromodomain-containing proteins (BRDs) sufficiently to account for their phenotypic effects. We describe highly specific inhibitors of ERK5 that do not inhibit BRDs. With these, we show that cellular inflammation and proliferation are not dependent on ERK5 catalytic activity, thus making ERK5 unique among the MAP kinases.

## Introduction

Extracellular signal-regulated kinase 5 (ERK5, BMK1) is a member of the mitogen-activated protein kinase (MAPK) family, which includes ERK1/2, JNK1/2/3, and p38α/β/δ/γ(1). However, unlike the other MAPK members, ERK5 contains a unique 400 amino acid C-terminal domain in addition to the kinase domain. Through the MAPK signaling cascade, MEK5 has been shown to activate ERK5 by phosphorylation at the TEY motif in the N-terminal activation loop (2). This event unlocks the N- and C-terminal halves, allowing ERK5 to auto-phosphorylate multiple sites in its C-terminal region, which can then regulate nuclear shuttling and gene transcription (3, 4). Non-conical pathways (including cyclin dependent kinases during mitosis and ERK1/2 during growth factor stimulation) also exist for phosphorylation of sites in the ERK5 tail (5–7). While ERK5 has been demonstrated to directly phosphorylate transcription factors (8–10), the non-catalytic C-terminal tail of ERK5 can also interact with transcription factors and influence gene expression (4, 11, 12).

ERK5 can be activated in response to a range of mitogenic stimuli (e.g., growth factors, GPCR agonists, cytokines) and cellular stresses (e.g., hypoxia, shear stress) (13). Like most kinases including MAPK members, ERK5 function is assumed to be driven by its kinase activity. ERK5 deletion is embryonic lethal in mice and a variety of tissue- or development-stage restricted knockouts have shown clear phenotypes, suggesting that the catalytic function and/or an aspect of the non-kinase domain(s) have key roles in development and mature organ function (14–18). The availability of the first ERK5 inhibitor XMD8-92 enabled the study of phenotypes resulting from direct kinase inhibition (19, 20). Effects of this inhibitor on cell proliferation were shown to be comparable to overexpression of a dominant negative ERK5 mutant (20, 21) or to siRNA-mediated ERK5 knockdown (21, 22), implicating a key role of the kinase function. Multiple reports have shown promising and corroborating effects of ERK5 knockdown and pharmacological inhibition in controlling inflammation and tumor growth (20, 22–26).

Given the proposed therapeutic uses for ERK5 inhibitors, we set out to develop improved compounds with high selectivity and potency. With these new ERK5 compounds, we show that first generation ERK5 inhibitors actually derived the bulk of their biological activity from off-target activity on bromodomains (BRDs). Surprisingly, selective inhibition of the kinase activity alone had no effect on cellular immune response or proliferation, in contrast to whole ERK5 protein knockdown. The newly developed ERK selective inhibitors should prove useful in determining which phenotypes derive from catalytic activity and which derive from other functions of this multi-domain kinase.

## Results and Discussion

### Inhibitors of ERK5

We synthesized derivatives of the benzopyrimidodiazepinone XMD8-92, a first generation ERK5 kinase inhibitor (**Fig. 1**). Among these compounds were ATP-competitive inhibitors that potently inhibit ERK5 with IC_50_ values ranging from 8 to 190 nM using the chemoproteomics platform KiNativ, which profiles global kinase inhibition in complex lysates (27, 28). At a 1 μM screening concentration, there was no significant inhibition of any off-target kinases among the greater than 100 kinases profiled in a single cellular lysate, indicating excellent specificity (**Supplementary Data S1**).

**Fig. 1.**
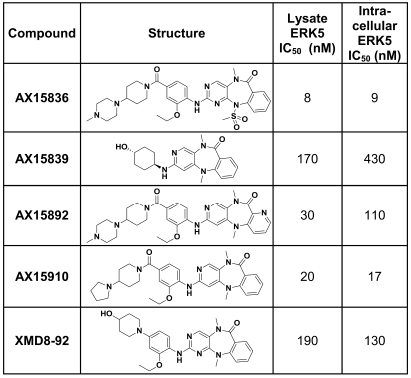
**Compound structures and potencies against ERK5 in cell lysate and in live cells treated with compound.**

The ERK5 inhibitors were also evaluated in an additional KiNativ experimental format wherein compound is incubated with live cells prior to washing, lysis and analysis. Such experiments provide an indication of the cellular permeability of the compound and of intracellular target- (and off target-) engagement. As seen in **Fig. 1**, intracellular ERK5 IC_50_ values were very similar to lysate ERK5 IC_50_ values, demonstrating that the compounds effectively reached their intracellular target. The most selective ERK5 inhibitor AX15836 was also profiled using live cell KiNativ across several cell types, including PBMCs, endothelial cells, and oncogenic cell lines, and verified to maintain its intracellular potency (4-9 nM) across all cells tested.

The compounds were next tested for their ability to inhibit epidermal growth factor (EGF)-mediated auto-phosphorylation of endogenous ERK5 in HeLa cells (4). HeLa cells are commonly used to study ERK5 regulation in part due to the ability to clearly observe ERK5 activation resulting from triggers such as growth factors and mitosis (5, 29, 30). In this assay, activated ERK5 migrates more slowly than unactivated ERK5 on SDS-PAGE by virtue of the protein being phosphorylated. As seen in **Fig. 2**, when treated at 2 μM, a concentration of almost 5-fold over the weakest intracellular ERK5 IC_50_, all of the ERK5 inhibitors substantially blocked the formation of phosphorylated ERK5 upon EGF stimulation. Because ERK5 is an intracellular target, this provided additional evidence, together with the KiNativ results, that the compounds were able to effectively engage their intracellular target.

**Fig. 2.**
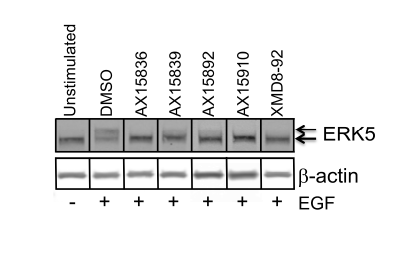
HeLa cell ERK5 auto-phosphorylation assay. Stimulation of HeLa cells with EGF induced a slower migrating, auto-phosphorylated ERK5 band (upper arrow) separated by SDS-PAGE and detected by western blot. This p-ERK5 was prevented by pharmacological ERK5 inhibition. Compounds were screened at 2 μM. Figure is a composite of lanes from nonadjacent samples on the same blot.

### Characterization of inhibitors in cellular models of inflammation

ERK5 has been recently studied as a target for mediating inflammation (23, 24, 26). We determined the activity of the new ERK5 compounds in cellular assays of inflammatory response. To function in the recruitment of neutrophils and monocytes, the endothelial cell adhesion molecule E-selectin is rapidly synthesized in response to inflammatory stimulation (31). Primary human umbilical vein endothelial cells (HUVEC) were incubated with compounds prior to stimulation with the toll-like receptor (TLR1/2) agonist Pam_3_CSK_4_. Up-regulation of cell-surface E-selectin was quantified by flow cytometry (**Table 1**). Similar to that reported by Wilhelmsen and colleagues (23), we found the ERK5 inhibitor XMD8-92 to inhibit up to 38% of the E-selectin expression. Likewise, two other ERK5 inhibitors AX15839 and AX15910 were effective in reducing E-selectin. Surprisingly, two of the more potent ERK5 inhibitors (AX15836 and AX15892) were inactive in this assay, even at a concentration of compound at least 90-fold higher than the intracellular ERK5 IC_50_ (**Fig. 1**).

**Table 1.**
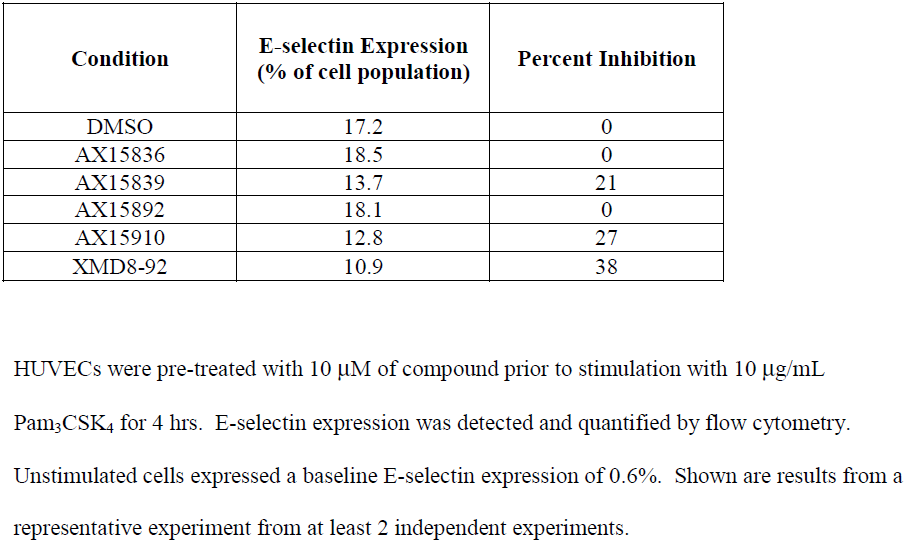
Surface E-selectin expression on endothelial cells stimulated with TLR1/2 agonist Pam_3_CSK_4_.

| Condition | E-selectin Expression<br>(% of cell population) | Percent Inhibition |
| --- | --- | --- |
| DMSO | 17.2 | 0 |
| AX15836 | 18.5 | 0 |
| AX15839 | 13.7 | 21 |
| AX15892 | 18.1 | 0 |
| AX15910 | 12.8 | 27 |
| XMD8-92 | 10.9 | 38 |

HUVECs were pre-treated with 10 μM of compound prior to stimulation with 10 $\mu$ g/mL
Pam<sub>3</sub>CSK<sub>4</sub> for 4 hrs. E-selectin expression was detected and quantified by flow cytometry.
Unstimulated cells expressed a baseline E-selectin expression of 0.6%. Shown are results from a representative experiment from at least 2 independent experiments.

Given these results, it seemed likely that an additional activity was responsible for the efficacy of AX15839, AX15910, and XMD8-92. We therefore performed a more extensive KiNativ experiment to expand the kinase coverage to more than 200 kinases and furthermore increased the compound screening concentration to 10 μM. However, we still did not find any significant shared inhibition of off-target kinases among these three inhibitors (**Supplementary Data S2**). XMD8-92 was derived from the polo-like kinase (PLK1) inhibitor BI-2536 (19, 20). Recently, BI-2536 (as well as a number of other kinase inhibitors) has been shown to inhibit the interaction between BRD and acetyl-lysine binding (32–34). BRDs are protein modules that bind to ε-N-acetylated lysine-containing sequences and modulate transcriptional processes. Members of the dual BRD-containing BET (bromo and extra terminal) proteins BRD2, BRD3, BRD4, and BRDT are targets of drugs currently pursued in oncology, neurological diseases, diabetes, atherosclerosis, and inflammation (35, 36). To determine whether XMD8-92 and other ERK5 kinase inhibitors can likewise inhibit BRDs, we screened the compounds against BRD4, the most well-studied and key member of the BET protein family.

**Table 2** shows the dissociation constants for compound binding to the first BRD of BRD4 ((BRD4(1)). Using the BROMOscan assay (DiscoveRx Corp., San Diego, CA), two reference BRD inhibitors, JQ1 (37) and I-BET762 (38) exhibited potent BRD4(1) K_d_ values. However, analysis of the ERK5 inhibitors using this method revealed a clear split. Compounds that were active in the E-selectin assay, AX15839, AX15910 and XMD8-92, potently interfered with the acetyl-lysine/ BRD4(1) interaction. These compounds thus represent dual inhibitors of ERK5 kinase and of BRD. In contrast, compounds that were potent on ERK5 but inactive in the E-selectin assay, AX15836 and AX15892, gave considerably higher BRD4(1) dissociation constants, indicating loss of off-target BRD inhibition.

**Table 2.** Inhibitor characteristics and classification.

| **Compound** | **ERK5 IC<sub>50</sub>**<br>**(nM)<sup>a</sup>** | **BRD4(1) K<sub>d</sub>**<br>**(nM)<sup>b</sup>** | **Ratio of ERK5**<br>**IC<sub>50</sub>:BRD4(1) K<sub>d</sub>** | **Inhibitor**<br>**Classification** |
| --- | --- | --- | --- | --- |
| AX15836 | 8 | 3600 | 0.002 | ERK5 |
| AX15892 | 30 | 610 | 0.049 | ERK5 |
| AX15839 | 170 | 130 | 1.3 | Dual |
| AX15910 | 20 | 22 | 0.9 | Dual |
| XMD8-92 | 190 | 170 | 1.1 | Dual |
| I-BET762 | >10000 | 31 | >320 | BRD |
| JQ1 | >10000 | 6 | >1670 | BRD |

<sup>a</sup>ERK5 IC<sub>50</sub> value determined using KiNativ
<sup>b</sup>BRD4(1) K<sub>d</sub> value determined at DiscoveRx

Knowing that the dual ERK5/BRD inhibitors were efficacious in the E-selectin HUVEC assay whereas the ERK5-selective inhibitors had no effect (**Table 1**), we returned to that assay to measure the activity of the two BRD-selective reference inhibitors. Using 1 μM of I-BET762 and JQ1, we observed E-selectin reductions of 27 and 29%, respectively, confirming the notion that TLR1/2-induced E-selectin expression in endothelial cells could be reduced by BRD inhibition but not by ERK5 inhibition. Neither BRD inhibitor was active against ~100 kinases (including ERK5) when profiled at up to 10 μM using KiNativ (**Supplementary Data S3**).

We characterized the activities of several compounds from each classification type in additional cellular models of inflammation and found good consistency of response. For brevity, we show the results of three representative compounds: AX15836 as the ERK5-selective inhibitor, AX15839 as the dual ERK5/BRD inhibitor, and I-BET762 as the selective BRD inhibitor.

To determine whether the compounds could suppress inflammatory cytokine response, endothelial cells were pre-treated with compound and stimulated with Pam_3_CSK_4_. Culture supernatants were subjected to immunoassay for cytokines IL-6 and IL-8. As seen in **Table 3**, compounds with BRD inhibition (selective and dual) suppressed IL-6 and IL-8; however, the ERK5-specific inhibitor AX15836 was completely ineffective (EC_50_≫10μM), suggesting that it was the BRD inhibition component of the compounds that mediated cytokine reduction.

**Table 3.** EC_50_ values of compounds in reducing cytokines IL-6 and IL-8 produced by endothelial cells stimulated with TLR1/2 agonist Pam_3_CSK_4_.

| Compound | Inhibitor Classification | EC<sub>50</sub> (μM) of cytokine reduction |  |
| --- | --- | --- | --- |
|  |  | IL-6 | IL-8 |
| AX15836 | ERK5 | >>10 | >>10 |
| AX15839 | Dual | 1.73 | 1.79 |
| I-BET762 | BRD | 0.15 | 0.12 |

HUVECs were pre-treated with compound prior to stimulation with 10 μg/mL Pam<sub>3</sub>CSK<sub>4</sub> for 48 hrs. Culture supernatants were tested for cytokines by immunoassay. Shown are results from a representative experiment from at least 2 independent experiments.

To determine whether the lack of ERK5-specific effect was limited to a certain cell type and agonist, we repeated the experiment using a normal human bronchial epithelial cell line, BEAS-2B. ERK5 has been identified to be part of an IL-17-mediated signaling cascade that drives keratinocyte proliferation and tumorigenesis (39). IL-17A is also overexpressed in conditions of chronic inflammation such as asthma and is thought to mediate airway neutrophilia through the induction of additional cytokines from target cells (40–42). After pre-incubation with compound, BEAS-2B cells were stimulated with the pro-inflammatory cytokine IL-17A. IL-6 and IL-8 cytokine release from the bronchial epithelial cells were measured by immunoassay (**Table 4**). Again, the ERK5-selective compound AX15836 had no effect on these induced cytokines. In contrast, inhibitors with BRD inhibition activity suppressed inflammatory cytokine response to IL-17A in this cell type.

**Table 4.**
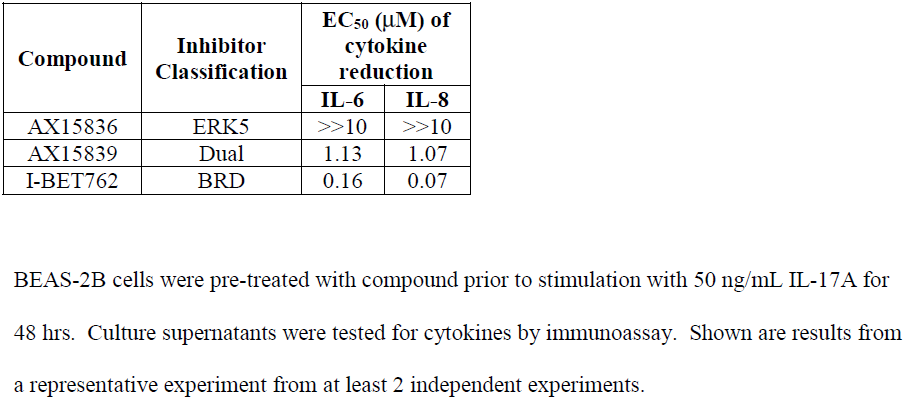
EC_50_ values of compounds in reducing cytokines IL-6 and IL-8 produced by bronchial epithelial cells stimulated with IL-17A.

| Compound | Inhibitor Classification | EC<sub>50</sub> (μM) of cytokine reduction |  |
| --- | --- | --- | --- |
|  |  | IL-6 | IL-8 |
| AX15836 | ERK5 | >>10 | >>10 |
| AX15839 | Dual | 1.13 | 1.07 |
| I-BET762 | BRD | 0.16 | 0.07 |

BEAS-2B cells were pre-treated with compound prior to stimulation with 50 ng/mL IL-17A for 48 hrs. Culture supernatants were tested for cytokines by immunoassay. Shown are results from a representative experiment from at least 2 independent experiments.

Cellular function of a protein is often studied by reducing its expression via RNA interference. The interpretability of removing the entire protein is typically justified when small molecule inhibitors can demonstrate the same phenotype. In multiple studies (20, 22–25), ERK5 kinase inhibition using XMD8-92 has been employed in parallel to siRNA-mediated knockdown of ERK5 to show two lines of supporting evidence for ERK5’s role. Given the lack of cellular effect by selective ERK5 kinase inhibitors, we used siRNA to deplete ERK5 in the HUVEC and BEAS-2B cells to evaluate the role of ERK5 presence. Cells were transfected with siRNA to significantly reduce ERK5 protein expression, as confirmed by western blots on days 2 and 4 post-transfection (**Fig. 3 *A*, *B***). On day 2 post-transfection, cells were stimulated with the respective agonists, and the culture supernatant was collected 2 days later (day 4 post-transfection) for cytokine analysis. Indeed, depletion of ERK5 protein resulted in a reduction in IL-6 and IL-8 in both endothelial and epithelial cells (**Fig. 3 *C*, *D***). These data show that the entire ERK5 protein is necessary for modulating inflammatory response. The contrasting lack of effect by specific small molecule-mediated inhibition of ERK5 kinase activity suggests that a non-catalytic function of ERK5 plays a more important role.

**Fig. 3.**
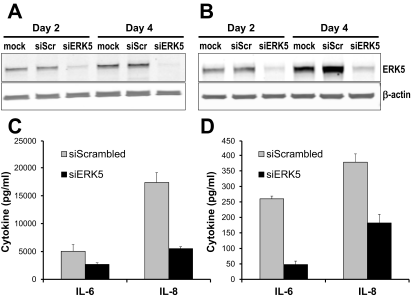
ERK5 protein depletion by siRNA suppresses inflammatory cytokine response. HUVEC **(*A*)** and BEAS-2B **(*B*)** cells were transiently transfected with mock treatment, scrambled siRNA control (siScr), or siRNA against ERK5 (siERK5). Expression of ERK5 and β-actin was monitored by western blot on day 2 (addition of Pam3CysK4 or IL-17) and day 4 (collection of culture supernatant for cytokine analyses). Immunoassay of IL-6 and IL-8 from treated HUVEC **(*C*)** and BEAS-2B **(*D*)** cells indicates significant suppression of inflammatory cytokines when ERK5 is depleted. Figures are representative experiments from 2 independent experiments.

We searched for an anti-inflammatory effect of AX15836 in additional cellular models of innate and adaptive immunity from both murine and human sources, as listed in **Supplementary Table S1**. Inhibition of BRD/acetyl-lysine binding with either JQ1 or I-BET762 resulted in efficacy in several models including primary cells such as PBMCs. In contrast, the ERK5-only inhibitor AX15836 was ineffective against all models tested. We therefore conclude that ERK5 kinase activity does not have a role in cellular immune response.

### Characterization of inhibitors in cancer cell proliferation and viability

ERK5 and BRDs have been separately studied as targets for oncology. ERK5 was proposed to control multiple processes important for tumorigenesis, including cellular proliferation, survival, invasion, and angiogenesis (43). To re-evaluate the role of ERK5 kinase inhibition in cancer cell proliferation, we tested our panel of inhibitors for effects on two cancer cell lines in which ERK5 has been characterized to mediate cell growth and survival. MM.1S multiple myeloma cells express ERK5, which can be activated by IL-6, a growth factor for this cancer type (44). IL-6-induced ERK5 activation was prevented by ERK5 inhibitors (specific ERK5 inhibitor AX15836 and dual ERK5/BRD inhibitor AX15839) (**Fig. 4*A***), confirming target engagement. Overexpression of a dominant negative form of ERK5 was reported to block IL-6-induced MM.1S proliferation (44). We compared IL-6-dependent proliferation in the presence or absence of the highly specific ERK5 inhibitor AX15836 (**Fig. 4*B***), but did not observe a significant effect on cell growth. As such, it seems likely that the reported anti-proliferation activity of overexpressing the dominant negative ERK5 mutant in MM.1S cells is the result of non-kinase-related or non-ERK5-related activity of that construct.

**Fig. 4.**
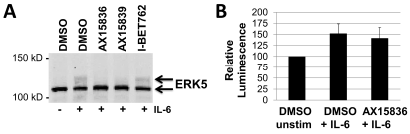
Multiple myeloma MM.1S cell growth is not affected by selective ERK5 inhibition. **(*A*)** IL-6 activates ERK5 in MM.1S cells, as shown by the appearance of a mobility retarded, auto-phosphorylated ERK5 band (upper arrow) separated by SDS-PAGE and detected by western blot. Inhibitors (2 μM) with ERK5 inhibitory activity (AX15836 and AX15839) prevented the induction of p-ERK5. **(*B*)** After 3 days of incubation, viable MM.1S cells were quantified in an assay measuring ATP content and expressed as relative luminescence. The ERK5-selective inhibitor AX15836 (1.67 μM) did not significantly reduce IL-6-dependent proliferation relative to the DMSO + IL-6 control. Data graphed in **(*B*)** are the mean ± SD of three independent experiments.

We likewise evaluated these inhibitors on the proliferation of the acute myeloid leukemia cell line MV4-11, which expresses the activating internal tandem duplication (ITD) mutation of FLT3 (FLT3-ITD). This driver mutation was reported to constitutively activate ERK5, and inhibition of the upstream kinase MEK5 led to reduced cell proliferation and viability (45). As previously noted in the literature (34), we found reference BRD inhibitors to be effective in this model, with EC_50_ values of 60 ± 10 nM and 170 ± 10 nM for JQ1 and I-BET762, respectively (mean ± SD of 3 experiments). Viability EC_50_s of the dual ERK5/BRD inhibitors (AX15839, AX15910, and XMD8-92) were less potent and ranged from 1.10 ± 0.25 to 3.28 ± 1.14 μM. Again, however, we observed no effect with the selective ERK5 compounds AX15836 and AX15892 (EC_50_s > 15 μM). Our studies thus demonstrate that highly specific pharmacological inhibition of ERK5 catalytic activity in cancer cell lines previously characterized to be regulated by this kinase had no effect on cell growth or viability in vitro. While xenograft studies might further delineate a more complex role of ERK5 kinase activity, pharmacokinetic characterization of AX15836 (**Supplemental Table S2**) did not indicate it to be optimal for in vivo dosing.

### Transcriptome analysis of cellular ERK5 kinase inhibition

The transcriptional role of ERK5 has previously been studied by overexpression of dominant negative mutants or by the use of inhibitors such as XMD8-92. We thus sought to analyze the effects of selective ERK5 inhibition in comparison to compounds with BRD inhibition (either dual ERK/BRD or BRD-only) on genome-wide gene expression. Two cellular models with reported ERK5-regulated signaling were used: Pam_3_CSK_4_-stimulated HUVECs (23, 24) as a model of inflammation, and EGF-stimulated HeLa cells (5, 20, 29) as an established cell model of ERK5 regulation. Cells were pre-incubated with DMSO vehicle, AX15836 (ERK5 inhibitor), AX15839 (dual ERK5/BRD inhibitor), or I-BET762 (BRD inhibitor), then stimulated with agonist. Cellular responses were verified by immunoassays and western blots using replicate wells in the same experiment.

RNA sequencing (RNA-Seq) of biological triplicates detected at least one transcript for 18925 genes in HUVEC samples and 17266 genes in HeLa samples. In both cell lines, samples treated with AX15836 showed very few genes to be differentially expressed (**Fig. 5*A***). The total number of genes using the default cut-off (adjusted p-value ≤ 0.1) for MA plots was 7 in HUVEC samples and 2 in HeLa samples. Moreover, the observed maximal fold-changes in expression when compared to the DMSO control samples were modest: below 1.6 and 2 for HUVEC and HeLa samples, respectively. Principal component analysis of all samples further confirmed the lack of differential gene expression in samples treated with the ERK5-only inhibitor AX15836. Conversely, cells treated with the dual ERK5/BRD inhibitor AX15839 and those treated with the BRD inhibitor I-BET762 showed a large number of differentially expressed genes (**Fig. 5*A***). The correlation of fold-changes in expression of those genes is shown in **Fig. 5*B***. The majority of the genes showed comparable expression patterns in that they were expressed at either higher or lower levels in each of the treated samples when compared to controls, confirming a shared regulatory mechanism between AX15839 and I-BET762.

**Fig. 5.**
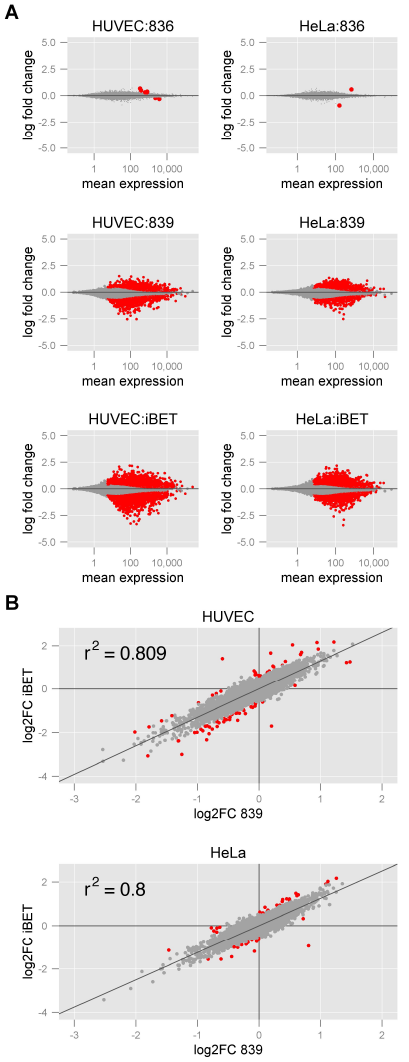
Differential gene expression in HUVEC and HeLa cells treated with AX15836, AX15839, and I-BET762 indicates lack of effect by selective ERK5 inhibition. **(*A*)** MA plots for HUVEC (left column) and HeLa cells (right column) treated with AX15836 (836, top row), AX15839 (839, middle row), and I-BET762 (iBET, bottom row). Differentially expressed genes with an adjusted p-value (DESeq2) of 0.1 or less are shown in red. **(*B*)** Correlation of gene expression profiles in HUVEC (top) and HeLa cells (bottom) treated with AX15839 and I-BET762. The log2-fold changes (log2FC) for compound-treated samples compared to the DMSO control samples are plotted for each gene. Outliers are highlighted in red and include those differentially expressed genes (at least a 1.5x fold-change and an adjusted p-value below 0.05 in one of the samples) with a residual outside of three times the standard deviation of all residuals.

Looking at individual genes of interest, AX15839 and I-BET762 significantly reduced Pam3CSK4-stimulated HUVEC gene expression of *IL6* (log2FC −0.72, *P* < 0.01 and log2FC −1.32, *P* < 0.001, respectively) and *CXCL8* (log2FC −0.73, *P* < 0.001 and log2FC −1.42, *P* < 0.001, respectively), consistent with the observed reductions in IL-6 and IL-8 proteins. *SELE* (E-selectin) transcripts were also reduced by these compounds (log2FC −0.47, *P* < 0.001 and log2FC −0.69, *P* < 0.001, respectively), consistent with the observed reduction in protein expression by flow cytometry. Additionally, both compounds with BRD inhibition (AX15839 and I-BET762) significantly suppressed transcription of other genes involved in inflammation, such as *IL7R* (IL-7 receptor) (log2FC −1.84, *P* < 0.001 and log2FC −2.38, *P* < 0.001, respectively), *PTGS2* (COX-2) (log2FC −.11, *P* < 0.001 and log2FC −1.65, *P* < 0.001, respectively), *CSF2* (GM-CSF) (log2FC −1.02, *P* < 0.001 and log2FC −1.60, *P* < 0.001, respectively), and *CCL2* (MCP-1) (log2FC −0.48, *P* < 0.001 and log2FC −1.41, *P* < 0.001, respectively), whereas inhibition of ERK5 kinase alone (AX15836) had no effect. Thus pharmacological inhibition of ERK5 kinase activity was not able to reduce inflammatory gene expression in endothelial cells, further supporting the concept that the previously-observed efficacy in first generation ERK5 inhibitors was due to an unrecognized inhibition of BRD/acetyl-lysine interaction.

We had shown that AX15836 could clearly inhibit the EGF-stimulated, phosphorylated form of ERK5 in HeLa cells, a frequently studied cell model of ERK5 regulation. We thus postulated that if the subsequent transcriptional effects of inhibiting ERK5 catalytic function could be seen, it would be in these cells. Still, we found no significant impact of AX15836 treatment. In contrast, the four genes most highly suppressed by both AX15839 and I-BET762 were: *HAS2* (hyaluronan synthase 2)(log2FC −2.53, *P* < 0.001 and log2FC −3.50, *P* < 0.001, respectively), *IL7R* (log2FC −2.08, *P* < 0.001 and log2FC −2.93, *P* < 0.001, respectively), *CXCL8* (log2FC −1.90, *P* < 0.001 and log2FC −2.14, *P* < 0.001, respectively), and *IL6* (log2FC −1.73, *P* < 0.001 and log2FC −2.69, *P* < 0.001, respectively). The transcription of both *HAS2* and *IL7R* have recently been reported to be potently downregulated by BET BRD inhibition in tumor cell lines and are thought to play key roles in cell growth and survival (46, 47). Consistent with previous observations that BRD inhibitors have differential effects on *MYC*, we also did not observe a reduction of *MYC* in HeLa cells (48); however, transcripts for cytokines IL-6 and IL-8, known to be increased in HeLa cells by EGF-mediated signaling (49), were suppressed by BRD inhibition. Our transcriptome profiling indicates that pharmacological inhibition of ERK5 kinase activity has no significant impact on gene transcription in two cell models, suggesting that phenotypes observed from genetic ablation of ERK5 result from kinase signaling-independent mechanisms.

## Conclusion

The biological function of ERK5 has been studied by a wide variety of experimental techniques. Genetic approaches in intact animals established a role for ERK5 in vascular and neuronal function. Depletion by siRNA in cell models implicated ERK5 in inflammatory and oncogenic pathways, and overexpression of a kinase-dead ERK5 mutant blocked numerous signaling events. Taken together, these experiments were useful to indicate the range of biological processes that ERK5 could influence. Since ERK5 is a kinase that has been demonstrated to phosphorylate transcription factors, the genetic phenotypes have been interpreted as being the result of removing the catalytic activity. However, the selective ERK5 inhibitors described here lack the anti-inflammatory and anti-proliferative effects induced by genetic manipulation of ERK5. The simplest explanation for this is that the immune and proliferation effects seen in ERK5 genetic models are due to ablation of non-kinase-related functions of ERK5. Separately, we demonstrate that previously reported ERK5 inhibitors, exemplified by XMD8-92, have off-target activity on an unrelated class of proteins, the BRDs. Our experiments show that BRD inhibition is sufficient to account for the anti-inflammatory and anti-proliferative cellular responses previously ascribed to ERK5 inhibition via XMD8-92. It appears to be pure happenstance that there is overlap between the phenotypes of BRD inhibition and ERK5 depletion/deletion.

Our findings show that ERK5 is a highly unusual MAP kinase. Like the other MAPK members, ERK5 is involved in a number of central biological pathways. In stark contrast however, the kinase activity of ERK5 appears to be unnecessary for many of its most widely studied functions.

## Materials and Methods

For determination of native kinase engagement, compounds were profiled against cell lysates or live cells using the KiNativ™ chemoproteomics platform (ActivX Biosciences, La Jolla, CA) (27, 28, 50). For flow cytometry, ERK5 auto-phosphorylation, immunoanalyses, and RNA-Seq, cells were pre-treated for an hour with compounds prior to agonist stimulation, followed by standard protocols. Experimental details and the synthetic schemes for AX15836, AX15839, AX15910, and AX15892 are provided in online **Supplementary Methods**.

## Acknowledgments

We express gratitude to our colleague and friend, the late Dr. Kevin Shreder for his invaluable contributions to this work. We thank Kai Nakamura, Lan Pham, Ray Li, and Julia Ayers for chemical syntheses, and Maria Sykes and Heidi Brown for their assistance on live cell KiNativ.

## Author Contributions

E.C.K.L., C.M.A., and J.I. developed and performed cell-based assays; T.K.N. planned and analyzed KiNativ studies. H.W. analyzed the RNA-Seq results. Y.H., S.S., and B.L. designed and synthesized inhibitors. E.C.K.L, T.K.N., H.W., Y.H., J.W.K., and J.S.R. wrote the paper.

## References

1. Morrison DK (2012) MAP kinase pathways. Cold Spring Harb Perspect Biol 4(11):a011254.

2. Mody N, Campbell DG, Morrice N, Peggie M, Cohen P (2003) An analysis of the phosphorylation and activation of extracellular-signal-regulated protein kinase 5 (ERK5) by mitogen-activated protein kinase kinase 5 (MKK5) in vitro. Biochem J 372(2):567–575

3. Kondoh K, Terasawa K, Morimoto H, Nishida E (2006) Regulation of nuclear translocation of extracellular signal-regulated kinase 5 by active nuclear import and export mechanisms. Mol Cell Biol 26(5):1679–1690

4. Morimoto H, Kondoh K, Nishimoto S, Terasawa K, Nishida E (2007) Activation of a C-terminal transcriptional activation domain of ERK5 by autophosphorylation. J Biol Chem 282(49):35449–35456

5. Diaz-Rodriguez E, Pandiella A (2010) Multisite phosphorylation of Erk5 in mitosis. J Cell Sci 123(18):3146–3156

6. Iñesta-Vaquera F a., et al. (2010) Alternative ERK5 regulation by phosphorylation during the cell cycle. Cell Signal 22(12):1829–1837

7. Honda T, et al. (2015) Phosphorylation of ERK5 on Thr732 is associated with ERK5 nuclear localization and ERK5-dependent transcription. PLoS One 10(2):e0117914.

8. English JM (1998) Identification of substrates and regulators of the mitogen-activated protein kinase ERK5 using chimeric protein kinases. J Biol Chem 273(7):3854–3860

9. Kamakura S, Moriguchi T, Nishida E (1999) Activation of the protein kinase ERK5/BMK1 by receptor tyrosine kinases: identification and characterization of a signaling pathway to the nucleus. J Biol Chem 274(37):26563–26571

10. Kato Y (1997) BMK1/ERK5 regulates serum-induced early gene expression through transcription factor MEF2C. EMBO J 16(23):7054–7066

11. Kasler HG, Victoria J, Duramad O, Winoto A (2000) ERK5 is a novel type of mitogen-activated protein kinase containing a transcriptional activation domain. Mol Cell Biol 20(22):8382–8389

12. Sohn SJ, Li D, Lee LK, Winoto A (2005) Transcriptional regulation of tissue-specific genes by the ERK5 mitogen-activated protein kinase. Mol Cell Biol 25(19):8553–8566

13. Nithianandarajah-Jones GN, Wilm B, Goldring CEP, Müller J, Cross MJ (2012) ERK5: Structure, regulation and function. Cell Signal 24(11):2187–2196

14. Yan L, et al. (2003) Knockout of ERK5 causes multiple defects in placental and embryonic development. BMC Dev Biol 3(1):11.

15. Regan CP, et al. (2002) Erk5 null mice display multiple extraembryonic vascular and embryonic cardiovascular defects. Proc Natl Acad Sci 99(14):9248–9253

16. Pan Y-W, et al. (2012) Inducible and conditional deletion of extracellular signal-regulated kinase 5 disrupts adult hippocampal neurogenesis. J Biol Chem 287(28):23306–23317

17. Hayashi M, et al. (2004) Targeted deletion of BMK1/ERK5 in adult mice perturbs vascular integrity and leads to endothelial failure. J Clin Invest 113(8):1138–1148

18. Angulo-Ibáñez M, et al. (2015) Erk5 contributes to maintaining the balance of cellular nucleotide levels and erythropoiesis. Cell Cycle 14(24):3864–3876

19. Deng X, et al. (2011) Discovery of a benzo[e]pyrimido-[5,4-b][1,4]diazepin-6(11H)-one as a potent and selective inhibitor of big MAP kinase 1. ACS Med Chem Lett 2(3):195–200

20. Yang Q, et al. (2010) Pharmacological inhibition of BMK1 suppresses tumor growth through promyelocytic leukemia protein. Cancer Cell 18(3):258–267

21. Bera A, et al. (2014) A positive feedback loop involving Erk5 and Akt turns on mesangial cell proliferation in response to PDGF. AJP Cell Physiol 306(11):C1089–C1100.

22. Rovida E, et al. (2015) The mitogen-activated protein kinase ERK5 regulates the development and growth of hepatocellular carcinoma. Gut 64(9):1454–1465

23. Wilhelmsen K, Mesa KR, Lucero J, Xu F, Hellman J (2012) ERK5 protein promotes, whereas MEK1 protein differentially regulates, the toll-like receptor 2 protein-dependent activation of human endothelial cells and monocytes. J Biol Chem 287(32):26478–26494

24. Wilhelmsen K, et al. (2015) Extracellular signal-regulated kinase 5 promotes acute cellular and systemic inflammation. Sci Signal 8(391):ra86–ra86.

25. Song C, et al. (2015) Inhibition of BMK1 pathway suppresses cancer stem cells through BNIP3 and BNIP3L. Oncotarget 6(32):1–11

26. Finegan KG, et al. (2015) ERK5 is a critical mediator of inflammation-driven cancer. Cancer Res 75(4):742–753

27. Patricelli MP, et al. (2011) In situ kinase profiling reveals functionally relevant properties of native kinases. Chem Biol 18(6):699–710

28. Patricelli MP, et al. (2007) Functional enterrogation of the kinome using nucleotide acyl phosphates. Biochemistry 46(2):350–358

29. Kato Y, et al. (1998) Bmk1/Erk5 is required for cell proliferation induced by epidermal growth factor. Nature 395(6703):713–6

30. Cude K, et al. (2007) Regulation of the G2–M cell cycle progression by the ERK5–NFκB signaling pathway. J Cell Biol 177(2):253–264

31. Peebles, Jr. RS, Bochner BS, Schleimer RP (1995) Pharmacologic regulation of adhesion molecule function and expression. Inflammation: Mediators and Pathways, eds Ruffolo, Jr. RR, Hollinger MA (CRC Press, Inc., New York).

32. Martin MP, Olesen SH, Georg GI, Schönbrunn E (2013) Cyclin-dependent kinase inhibitor dinaciclib interacts with the acetyl-lysine recognition site of bromodomains. ACS Chem Biol 8(11):2360–2365

33. Ember SWJ, et al. (2014) Acetyl-lysine binding site of bromodomain-containing protein 4 (BRD4) interacts with diverse kinase inhibitors. ACS Chem Biol 9(5):305–1171.

34. Ciceri P, et al. (2014) Dual kinase-bromodomain inhibitors for rationally designed polypharmacology. Nat Chem Biol 10(4):305–312.

35. Filippakopoulos P, Knapp S (2014) Targeting bromodomains: epigenetic readers of lysine acetylation. Nat Rev Drug Discov 13(5):337–56.

36. Padmanabhan B, Mathur S, Manjula R, Tripathi S (2016) Bromodomain and extra-terminal (BET) family proteins: New therapeutic targets in major diseases. J Biosci 41(2):295–311.

37. Filippakopoulos P, et al. (2010) Selective inhibition of BET bromodomains. Nature 468(7327):1067–1073.

38. Mirguet O, et al. (2013) Discovery of epigenetic regulator I-BET762: lead optimization to afford a clinical candidate inhibitor of the BET bromodomains. J Med Chem 56(19):7501–7515.

39. Wu L, et al. (2015) A novel IL-17 signaling pathway controlling keratinocyte proliferation and tumorigenesis via the TRAF4-ERK5 axis. J Exp Med 212(10):1571–1587

40. Laan M, Lötvall J, Chung KF, Lindén A (2001) IL-17-induced cytokine release in human bronchial epithelial cells in vitro☐: role of mitogen-activated protein (MAP) kinases. Br J Pharmacol 133(1):200–206.

41. Chesné J, et al. (2014) IL-17 in severe asthma. Where do we stand? Am J Respir Crit Care Med 190(10):1094–1101.

42. Kawaguchi M, et al. (2001) Modulation of bronchial epithelial cells by IL-17. J Allergy Clin Immunol 108(5):804–809.

43. Lochhead PA, Gilley R, Cook SJ (2012) ERK5 and its role in tumour development. Biochem Soc Trans 40(1):251–256.

44. Carvajal-Vergara X, et al. (2005) Multifunctional role of Erk5 in multiple myeloma. Blood 105(11):4492–4499

45. Razumovskaya E, Sun J, Rönnstrand L (2011) Inhibition of MEK5 by BIX02188 induces apoptosis in cells expressing the oncogenic mutant FLT3-ITD. Biochem Biophys Res Commun 412(2):307–312.

46. Lockwood WW, Zejnullahu K, Bradner JE, Varmus H (2012) Sensitivity of human lung adenocarcinoma cell lines to targeted inhibition of BET epigenetic signaling proteins. Proc Natl Acad Sci 109(47):19408–19413.

47. Ott CJ, et al. (2012) BET bromodomain inhibition targets both c-Myc and IL7R in high-risk acute lymphoblastic leukemia. Blood 120(14):2843–2852.

48. Fowler T, et al. (2014) Regulation of MYC expression and differential JQ1 sensitivity in cancer cells. PLoS One 9(1):e87003.

49. Amit I, et al. (2007) A module of negative feedback regulators defines growth factor signaling. Nat Genet 39(4):503–512.

50. Okerberg ES, et al. (2014) Monitoring native p38α: MK2/3 complexes via trans delivery of an ATP acyl phosphate probe. J Am Chem Soc 136(12):4664–4669

